# PDXGEM: Patient-Derived Tumor Xenograft based Gene Expression Model for Predicting Clinical Response to Anticancer Therapy in Cancer Patients

**DOI:** 10.1101/686667

**Authors:** Youngchul Kim, Daewon Kim, Biwei Cao, Rodrigo Carvajal, Minjung Kim

## Abstract

**Background:** Cancer is a highly heterogeneous disease and shows varying responses to anti-cancer drugs. Although several approaches have been attempted to predict the drug responses by analyzing molecular profiling data of tumors from preclinical cancer models or cancer patients, there is still a great need of developing highly accurate prediction models of response to the anti-cancer drugs for clinical applications toward personalized medicine. Here, we present PDXGEM pipeline to build a predictive gene expression model (GEM) for cancer patients’ drug responses on the basis of data on gene expression and drug activity in patient-derived xenograft (PDX) models.

**Results:** Drug sensitivity biomarkers were identified by an association analysis between gene expression levels and post-treatment tumor volume changes in PDX models. Only biomarkers with concordant co-expression patterns between the PDX and cancer patient tumors were used in random-forest algorithm to build a drug response prediction model, so called PDXGEM. We applied PDXGEM to several cytotoxic chemotherapy as well as targeted therapy agents that are used to treat breast cancer, pancreatic cancer, colorectal cancer, or non-small cell lung cancer. Significantly accurate predictions of PDXGEM for pathological and survival outcomes were observed through extensive validation analyses of multiple independent cancer patient datasets obtained from retrospective observational study and prospective clinical trials.

**Conclusion:** Our results demonstrated a strong potential of utilizing molecular profiles and drug activity data of PDX tumors for developing a clinically translatable predictive cancer biomarkers for cancer patients. PDXGEM web application is publicly available at http://pdxgem.moffitt.org.

## Background

Cytotoxic chemotherapy and targeted therapy play important roles with surgery, radiotherapy, as well as a recent breakthrough immunotherapy in the treatment of cancer. Responses of cancer patients to drugs of those anticancer therapies largely vary because of the substantial heterogeneity in molecular characteristics of their tumors even with a histologically same type of cancer^1^. Although a considerable number of novel anticancer drugs have been introduced during the past few decades, overall survival and quality of life of cancer patients has not been improved much because mainly of unselective use of them in the presence of such heterogeneity in patients’ tumor characteristics and responses to the drugs^2^. Hence, it is necessary to develop a personalized anticancer therapy that can guide individual patients with heterogeneous tumors to be treated with mostly beneficial anticancer drugs. A successful personalized anticancer therapy will then greatly depend on predictive cancer biomarkers that can accurately select patients who will receive a benefit from the anticancer drugs^3^.

For a predictive cancer biomarker discovery, it is considered most desirable to analyze molecular profiling data and clinical outcomes of cancer patients obtained before or/and after a treatment with anticancer drugs of interest via a randomized clinical trial^4^. However, it is not straightforward to develop cancer biomarkers in this manner due to extremely huge cost and time spent in the process of the clinical trial. Especially, it is more challenging when developing cancer biomarkers for a single drug or a drug combination that were once a standard of care, but no more after a new drug introduction. Hence, many cancer biomarker studies depend heavily on testing anticancer drugs in preclinical cancer models including immortalized cancer cell lines and animal models^5^.

Cancer cell lines cultured *in vitro* have been widely utilized to understand molecular characteristics and drug activity mechanism of tumor cells. For instance, two large cancer cell line panels, Genomics of Drug Sensitivity in Cancer and Cancer Cell Line Encyclopedia, were established to develop new anticancer drugs and identify new molecular drug targets and predictive biomarkers by interrogating pharmacogenomic mechanisms in more than 1,000 cancer cell lines^6,7^. We and many other research teams have been developing techniques to translate *in vitro* cancer cell lines-driven biomarkers into prediction of cancer patients’ anticancer drug responses^8-13^. However, it still lacks well-validated biomarkers and biomarker discovery methods mainly because cancer cell lines undergo frequent occurrences of genetic variants and uncontrollable contaminations during cell culture and thereby there exists a clear disconnection between *in vitro* cancer cell lines and human tumor cells^14^.

Recently, patient-derived tumor xenograft (PDX) on which surgically derived patient’s tumor is implanted has been recognized to better inform therapeutic development strategies than the cancer cell lines. Large PDX-based studies such as National Cancer Institute MicroXeno project, Novartis PDX panel, and EuroPDX consortium study interrogated molecular characteristics based on multiplex molecular platforms including gene expression and genetic mutation, and reported that PDXs can retain the distinct characteristics of different tumors from different patients and therefore effectively recapitulate the intra-tumor and inter-tumor heterogeneity that represents human cancer^15-18^. These novel and unprecedented PDX resources can provide a new great opportunity to discover highly predictive cancer biomarkers that can guide cancer patients to highly beneficial anticancer therapeutics and accelerate the process of new drug development. However, very few attempts have been made so far and no analytic tool for developing a PDX-based predictive gene expression model (GEM) is yet available. We thus develop a new pharmacogenomic pipeline, so-called PDXGEM, that can facilitate building a highly predictive GEM of clinical response of cancer patients to anti-cancer drug on the basis of gene expression profile and drug response screening data of the preclinical PDX tumors.

In this study, we provide a full description of the PDXGEM and illustrate its utility by applying it to several cytotoxic and targeted therapeutic agents, and validating the prediction performance of resultant multi-gene expression models on independent external cancer patient cohorts with well-annotated clinical outcomes. We also have created a publicly available web-based application with an initial inventory of the data of the Novartis PDX panel and cancer patient cohorts that were used for PDXGEM development and validation.

## Results

The PDXGEM pipeline consists of four subsequent steps, 1) drug sensitivity biomarker discovery, 2) concordant co-expression analysis (CCEA), 3) multi-gene expression model training for drug response prediction, and 4) model validation (Figure 1; see *Materials and Method*). To demonstrate the utility of the PDXGEM, we applied the PDXGEM for building predictive GEMs of cancer patients’ responses to each of three chemotherapy agents and three targeted therapy drugs: paclitaxel and trastuzumab for breast cancer, 5FU and cetuximab for colorectal cancer (CRC), gemcitabine for pancreatic cancer, and erlotinib in non-small cell lung cancer (NSCLC). External validations of the resultant GEMs were conducted using publicly available gene expression data and clinical outcome data of independent cancer patient cohorts from prospective clinical trials or observational studies.

**Figure 1.**
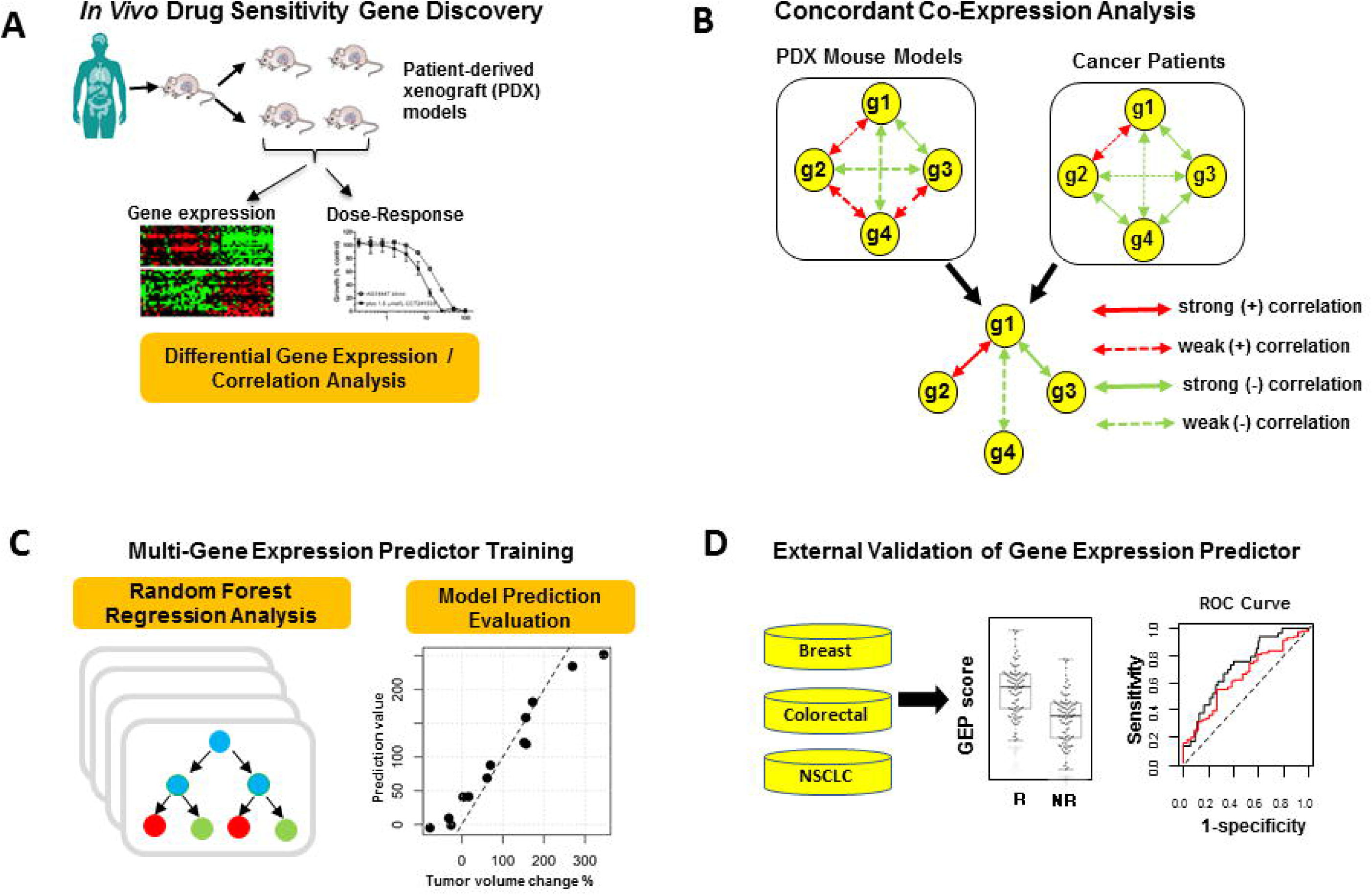
Schema of PDXGEM. a) In drug sensitivity gene discovery step, correlation analysis and differential expression analysis of gene expression data and drug-activity data in PDX tumors were conducted. (b) Concordant co-expression analysis selected drug sensitivity genes with concordant co-expression patterns between PDX tumors and cancer patients’ tumors (c) multi-gene expression model of drug response was trained using the random-forest algorithm (d) PDXGEM drug response prediction model was tested on independent cancer patient cohorts.

### PDXGEM for Predicting Paclitaxel Response in Breast Cancer Patients

Paclitaxel is a cornerstone of the current standard chemotherapy with fluorouracil, doxorubicin, and cyclophosphamide (FAC) for breast cancer treatment. PDXGEM was applied to build a multi-gene expression model to predict who may achieve a pathological complete response (pCR) to paclitaxel. 600 probesets were first identified as initial drug sensitivity biomarkers to exhibit differential expressions between three breast cancer PDXs with shrunken tumor volumes and ten breast cancer PDXs with increased tumor volumes after receiving paclitaxel (*t*-test p < 0.05, Figure 2A). The pattern of co-expression among the drug sensitivity genes, as measured by a gene-gene correlation coefficient, in the breast cancer PDXs were then quite distinct from that of breast cancer patients, in line with a previous study that there can be an inherent biological gap between PDX tumors and their origin cancer patient tumors because of different growth environments surrounding the tumors (Figure 2B; top). In the following concordant co-expression analysis (CCEA), 147 (24.5%) of the drug sensitivity biomarkers showed significant concordant co-expression coefficients (CCEC) from 0.204 to 0.464 between those breast cancer PDXs and a cohort of 251 breast cancer patients (GSE3494^19^), and we hereafter referred to as the concordant co-expression (CCE) biomarkers. The CCE biomarkers showed more concordant co-expression patterns across the breast cancer PDXs and patients with an increased median CCEC of 0.272, as compared to all drug sensitivity biomarkers with a median CCEC of 0.09 (Figure 2B; bottom). Next, a random forest (RF) predictor was trained using gene expression data of the breast cancer PDXs for all the CCE biomarkers as a model training set. A resultant RF predictor consisted of 145 CCE biomarkers possessing a positive variable importance value (Supplementary Fig. 1). In the training set, prediction scores of the RF predictor was tightly correlated with the observed tumor volume changes (r=0.982, p<0.01; Figure 2C).

**Figure 2.**
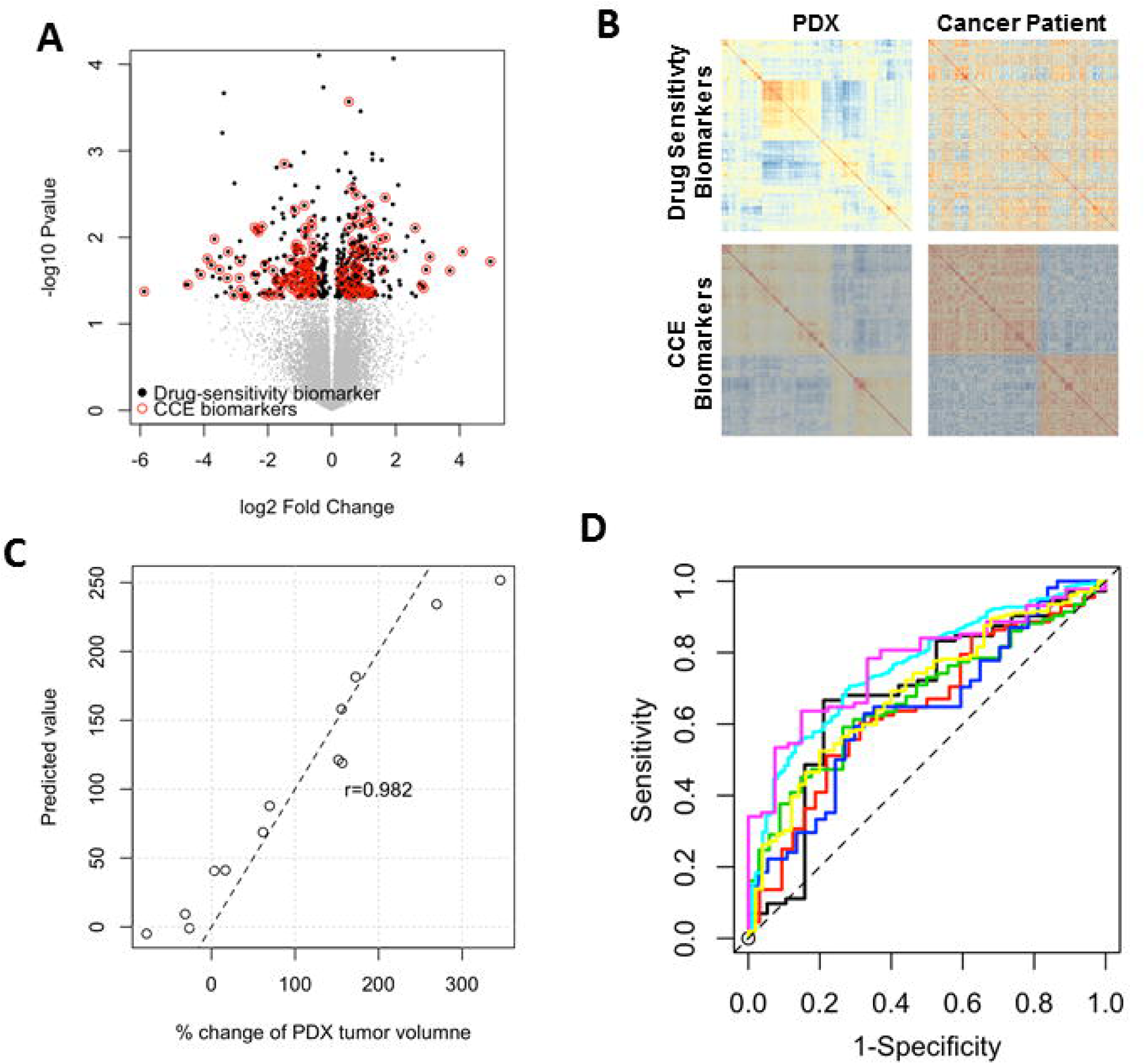
Development of PDXGEM for paclitaxel response prediction in breast cancer patient. (a) Volcano plot with log2 fold change of different gene expression levels (x-axis) in sensitivity and resistant PDX models and –log 10 p-value (y-axis). Black dots displays initial drug sensitivity biomarkers and red circle further indicates concordantly co-expressed biomarkers between PDX models and breast cancer patients. (c) Pearson correlation coefficient between observed percent change in PDX tumor volumes (x-axis) and PDXGEM prediction scores for breast cancer PDX models (y-axis) was 0.982. (d) ROC curves of paclitaxel PDXGEM on seven different breast cancer data sets.

To ensure the predictive performance of the RF predictor, we validated it with the use of seven independent gene expression datasets of breast cancer patients that were collected through four randomized clinical trials (GSE20271^20^, GSE22226^21^, GSE41998^22^, GSE42822^10^), two prospective observational studies (GSE25065^23^, GSE32646^24^), and one retrospective study cohort (GSE20194^25^). Notably, there were significant differences in prediction scores between patients with pCR and those with residual of disease (RD) after paclitaxel-based chemotherapy in all the breast cancer cohorts (p<0.05; Supplementary Fig. 2 A-G). In addition, area under the receiver-operating characteristic (ROC) curve (AUC) as an overall classification accuracy ranged from 0.653 to 0.789 (Figure 2D and Supplementary Fig. 2 A-G). To further determine whether the RF predictor is predictive for paclitaxel-specific response, we tested the predictor in 87 breast cancer patients who did not receive paclitaxel but only FAC combination chemotherapy in the GSE20271 clinical trial cohort. Consequently, there was no significant difference in prediction scores, suggesting that our predictor is predictive of response specifically to paclitaxel (AUC=0.589, p=0.44; Supplementary Fig. 2H).

To examine the utility of CCEA, we trained a RF predictor using all the 600 initial drug sensitivity biomarkers that did not undergo CCEA. Although this predictor was about three times complex as the above final RF predictor, there was no significant difference in its prediction scores between pCR and RD groups in four breast cancer patient and even decreased AUCs were observed in the remaining validation sets, suggesting that the CCEA could lead to a parsimonious gene expression model with more accurate prediction performance (Supplementary Figure 3).

**Figure 3.**
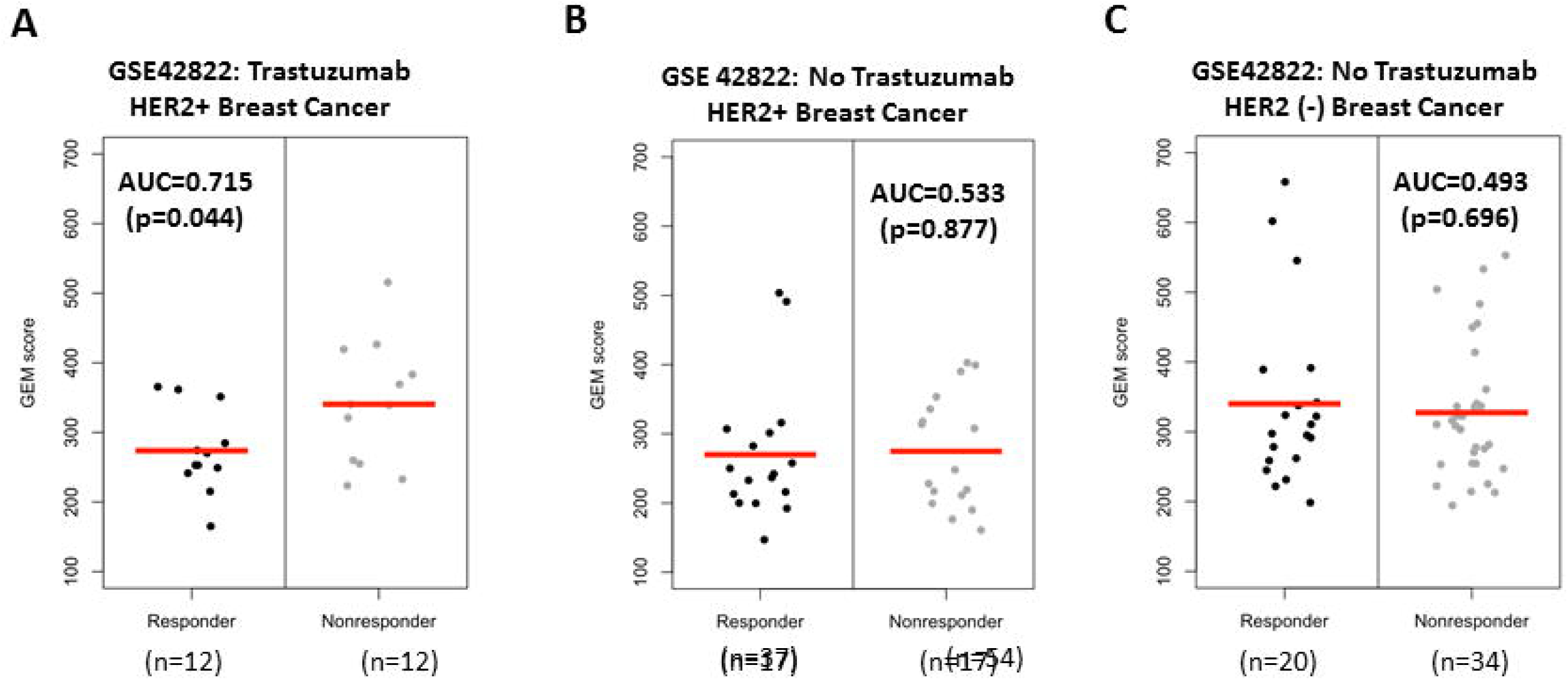
PDXGEM prediction scores for Trastuzumab in breast cancer patients by HER2 status. Distributional plot of PDXGEM prediction scores between responders and non-responders after receiving Transtuzumab (a) in HER2 positive breast cancer patients, (b) in HER2 positive breast cancer patients who did not receive Trastuzumab but other chemotherapy and (c) HER2 negative breast cancer patients who did not receive Trastuzumab. Red center lines represent the mean of prediction scores.

A gene ontology (GO) analysis for understanding the biological functions of 145 biomarkers of our final paclitaxel response predictor resulted in that COL1A1, RPH3AL, and THSD4 were the most significantly associated with breast neoplasm function (false discovery rate (FDR) p<0.001). In addition, DNA replication proteins and mismatch repair were top two representative pathways.

### PDXGEM for Trastuzumap-specific Response in Breast Cancer Patients

Trastuzumab is a monoclonal antibody used to treat human epidermal growth factor receptor 2 (HER2) positive breast cancer by itself or in combination with other anti-cancer therapeutics^26^. To construct a gene signature predicting response to the trastuzumab in breast cancer patients, we applied PDXGEM to data on gene expression profiles and post-treatment tumor volume changes in 13 breast cancer PDXs that underwent a monotherapy with trastuzumab. 1,333 drug sensitivity biomarkers were first identified with significant Spearman rank correlation relationships between gene expression levels and the tumor volume changes, and 515 CCE biomarkers were further screened out with significant CCEC in a range from 0.201 to 0.509. Finally, an optimal predictor was constructed with 480 CCE biomarkers possessing positive variable importance in RF model training analysis and the model yielded a strong correlation coefficient of 0.977 between predicted and observed tumor volume changes in the breast cancer PDX models. An independent validation of this RF predictor was performed using data from the US Oncology 02-103 breast cancer trial (GSE42822^10^) where 25 patients with stage II-III HER2-positive breast cancer received trastuzumab. We observed a borderline significant difference in prediction scores between 12 patients with pCR and 13 patients with RD after treatment with trastuzumab (AUC=0.712, p=0.074). Considering the large number of the biomarkers involved in the predictor and the encouraging AUC value, we set the more stringent threshold value of 0.3 for CCEC at the 2^nd^ step of PDXGEM to yield a less complex GEM with more concordantly co-expressed biomarkers between the breast cancer PDXs and patients. As expected, a new RF predictor was constructed with 193 CCE biomarkers and yielded a more significant difference in prediction scores between pCR and RD response groups in the breast cancer trial cohort (AUC=0.737; p=0.025) (Figure 3A). To assess the specificity of the RF predictor for trastuzumab, we validated the RF predictor on 34 HER2-positive and 54 HER2-negative breast cancer patients who did not receive trastuzumab in the same clinical trial. In both HER2 strata, we observed no difference in prediction scores between pCR and RD response groups (AUC=0.533, p=0.877 for the HER2 positive breast cancer; AUC=0.493, p=0.696 for the HER2 negative breast cancer; Figure 3B). When the predictor was further tested using other available breast cancer patient cohorts under not Trastuzumab but paclitaxel-based chemotherapy, none of the breast cancer cohorts did show any significant difference in prediction scores, strongly suggesting that this RF predictor is predictive of trastuzumab-specific response in breast cancer patients (Supplementary Fig. 4).

**Figure 4.**
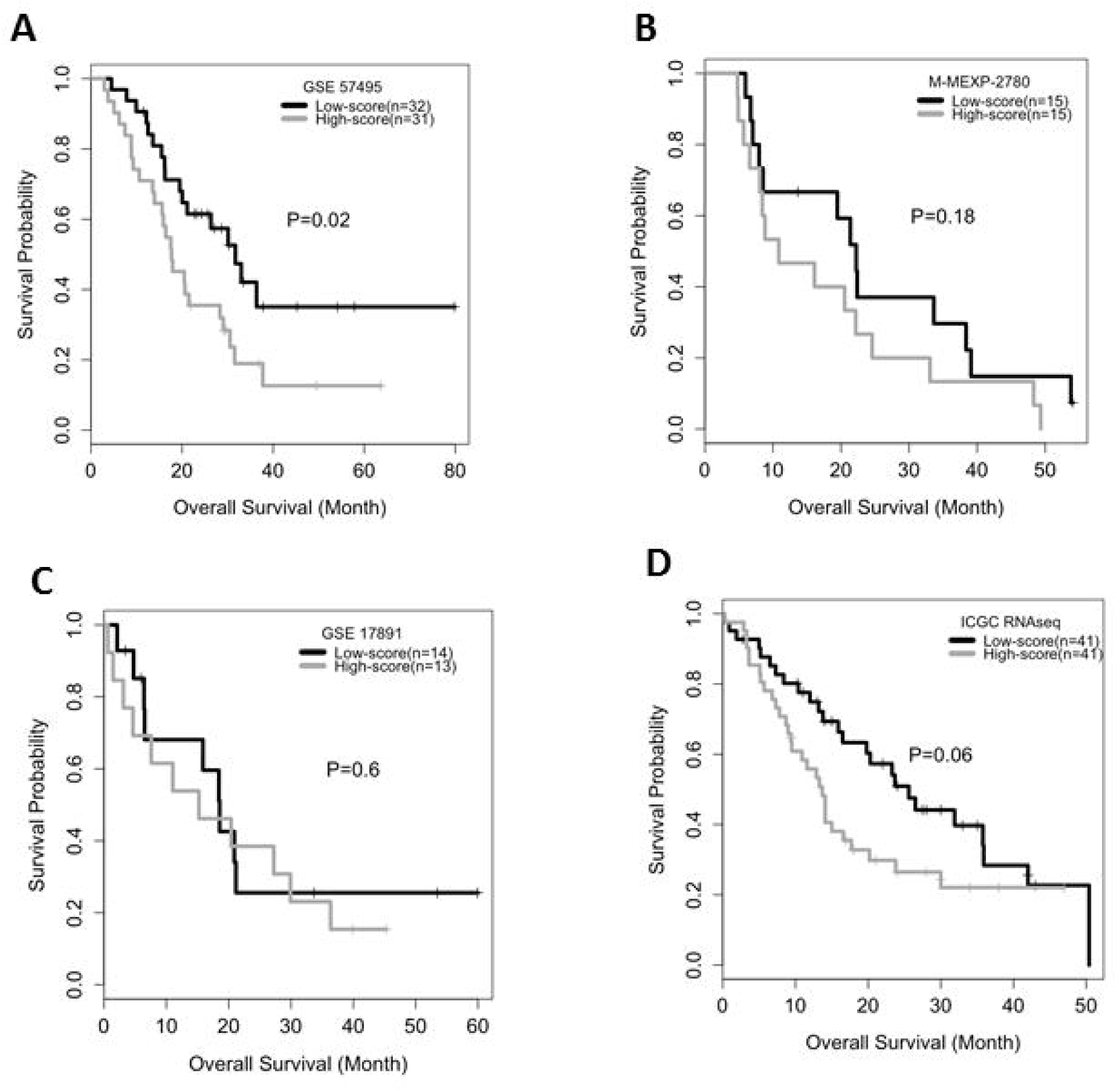
PDXGEM for Gemcitabine in Pancreatic Cancer. (a-d) Kaplan-Meier curves of overall survival between pancreatic cancer patients with a higher (gray) and lower (black) score than the median prediction score in GSE57495, M-MEXP-2789, GSE17891, and ICGC cohort. P-value was calculated using log-rank survival test.

Finally, a gene ontology analysis of the 193 biomarkers in the final predictor resulted in the most significant pathways including miRNA targets in ECM and membrane receptors, focal adhesion-PI3K-Akt-mTOR-signaling pathway, inflammatory response pathway, and apoptosis-related network due to altered Notch3 (FDR p<0.05). Especially, the PI3K-Akt-mTOR-signlaing pathway is a downstream pathway of HER2 and is well known to be responsible for promoting cell proliferation and angiogenesis^27^. In addition, COLTA1 gene had top second high variable importance in the RF model training analysis and it was reported in the genomic study in a phase 3 clinical trial for trastuzumab that COLTA1 is a key gene in integrin signaling pathway linked to a decreased recurrence-free survival time after adjuvant trastuzumab therapy^28^.

### PDXGEM for Predicting Response to Gemcitabine in Pancreatic Cancer Patients

Gemcitabine is currently used as backbone in a first line or second line treatment for pancreatic ductal adenocarcinoma (PDA) that carries a dismal prognosis with a typical overall survival (OS) of 6 months from diagnosis^29^. Although only six pancreatic cancer PDXs were available for tumor volume changes after receiving gemcitabine in the Novartis PDX panel, we attempted to use PDXGEM for developing a gene signature predicting response to gemcitabine.

965 drug sensitivity biomarkers were first screened out using *t*-test that contrasts expression levels of an individual probeset between two PDXs having shrunken tumor volumes and four PDXs having increased tumor volumes after receiving gemcitabine (p<0.05). 404 CCE biomarkers were further selected from CCEA using pretreatment gene expression data of 39 patients with PDA (GSE15471^30^). In a RF model training analysis of the PDX dataset, a final drug response prediction model was consisted of 298 CCE biomarkers with a high correlation coefficient of 0.959 between predicted scores and observed percent changes in tumor volumes.

In an external validation of predictive ability of this gene signature, we could collect gene expression data and survival outcome data from a retrospective study cohort of 63 patients with stage I/II PDA who received gemcitabine (GSE57495^31^). For a comparative analysis of the survival outcomes, we defined two patent subgroups according to whether patients’ prediction scores were higher or lower than the median prediction score. The low-score group then showed a significantly better OS (median OS=31.7 months, 95% CI=19.5∼not reached) than the high-score group (Figure 4A; median OS=7.7 months, 95% CI=13.5-28.3, log-rank p=0.023).

To assess the prediction ability of the signature for gemcitabine-specific response, we analyzed in the same manner a survival outcome data from a prospective observational study cohort of 30 patients with PDA who did not receive adjuvant chemotherapy (M-MEXP-2780, ArrayExpress). No significant difference was seen in OS, but the low-score group had a promising OS than the high-score group (Figure 4B; median OS=22.9 months for the low-score group, 10.9 months for the high-score group; log-rank p=0.18), implying that our PDXGEM signature could be not only predictive of gemcitabine response but also prognostic in part. To confirm the prognostic value of the predictor, we analyzed two additional cohorts of patients with PDA, 1) GSE17891^32^ (n=29) and 2) International Cancer Genome Consortium^33^ (ICGC, n=82) even though their chemotherapeutic treatment records were not available. For GSE17891 cohort, we again observed not significant but slightly better OS in the low-score group (p=0.6, Figure 4C). In addition, a multivariable Cox regression analysis revealed that higher prediction score was associated with higher risk of death (hazard ratio (HR)=1.087, 95% confidence interval (CI)=1.01-1.161, p=0.01), independent of known demographic and clinical prognostic factors of PDA including age at surgery, tumor stage, and molecular subtypes of PDA. For the ICGC cohort, there was a better OS significantly in the low-score group than the high-score group (log-rank test p=0.06; median OS=25.6 for the low-score group and 13.7 for the high-score group; Figure 4D), and the raw continuous prediction score was again significantly associated with OS (HR=1.026, 95%CI=1.001-1.051), independent of age and tumor stage. Although a further analysis will be needed in patient cohorts with data on detailed drug treatment records, these observations suggested that our PDXGEM predictor could be not only predictive of response to gemcitabine, but could also have a prognostic value in terms of predicting for a long-term outcome OS in patients with PDAC.

### PDXGEM for Predicting Response to 5FU in Colorectal Cancer Patients

5-fluorouracil (5FU) is widely used in the therapy of solid tumors, including colorectal, breast, head and neck cancers. Using PDXGEM, we built a gene signature predicting response to 5FU in colorectal cancer (CRC) patients by analyzing data of 16 colorectal cancer PDXs on gene expression and percent of change in tumor volumes after treatment with 5FU. At drug sensitivity biomarker discovery step, expression levels of 848 probesets were significantly correlated with the percent of change in tumor volumes (p<0.05). We next identified 332 CCE biomarkers from CCEA of the PDXs and a cohort of metastatic CRC patients (GSE14095^34^; n=189). In a following RF prediction training step, all the CCE biomarkers displayed positive variable importance and a resultant RF predictor yielded an almost perfect correlation coefficient of 0.978 between its prediction values and observed tumor volume changes in the PDX models. According to a gene ontology analysis of the biomarkers, the most significantly enriched function was amino acid catabolic process, which is in agreement with that 5-FU drug pathway is regulated via a complex network of anabolic and catabolic genes^35^.

As an external validation for a prediction performance of the RF predictor, we tested the predictor using two gene expression datasets of CRC patients. The first dataset (GSE62322^36^) was obtained from a phase 2 clinical trial where a percent of change in lesion size were assessed for 20 patients with liver metastatic CRC after receiving 5FU in combination with leucovorin and Irinotecan (FOLFIRI). Our RF predictor produced prediction scores with a significantly large difference between 9 responders and 11 non-responders (Figure 5A; AUC=0.788, 95% CI=0.56-0.99, p=0.035). The other validation dataset was collected from a retrospective study (GSE39582^37^) and consisted of two CRC patient cohorts: 1) 75 primary CRC patients treated with 5FU monotherapy, and 2) 69 primary and 20 metastatic CRC patients who received 5FU as either FOLFIRI or FOLFOX combination therapies^37^. We divided patients into three balanced groups (low-, intermediate-, and high-score groups) by breaking their prediction scores at tertiles of the prediction scores and examined a survival trend across the three groups. In the 5FU monotherapy cohort, there was a trend for longer OS in primary CRC patients with lower prediction scores, but not statistically significant, which might be due to a low event rates (trend test p=0.319) (Figure 5B). For the combination therapy cohort, the low-score group had significantly better OS than the intermediate- and high-score groups in the 20 metastatic CRC patients (Figure 5C; median OS=41, 22, and 20 months for high-, intermediate-, and low-score strata; p=0.03). However, a completely reversed survival trend was observed in the 69 primary CRC patients, reflecting a known fact that adjuvant FOLFIRI is not effective for resected primary cancer in contrast to metastatic disease^38,39^ (Figure 5D).

**Figure 5.**
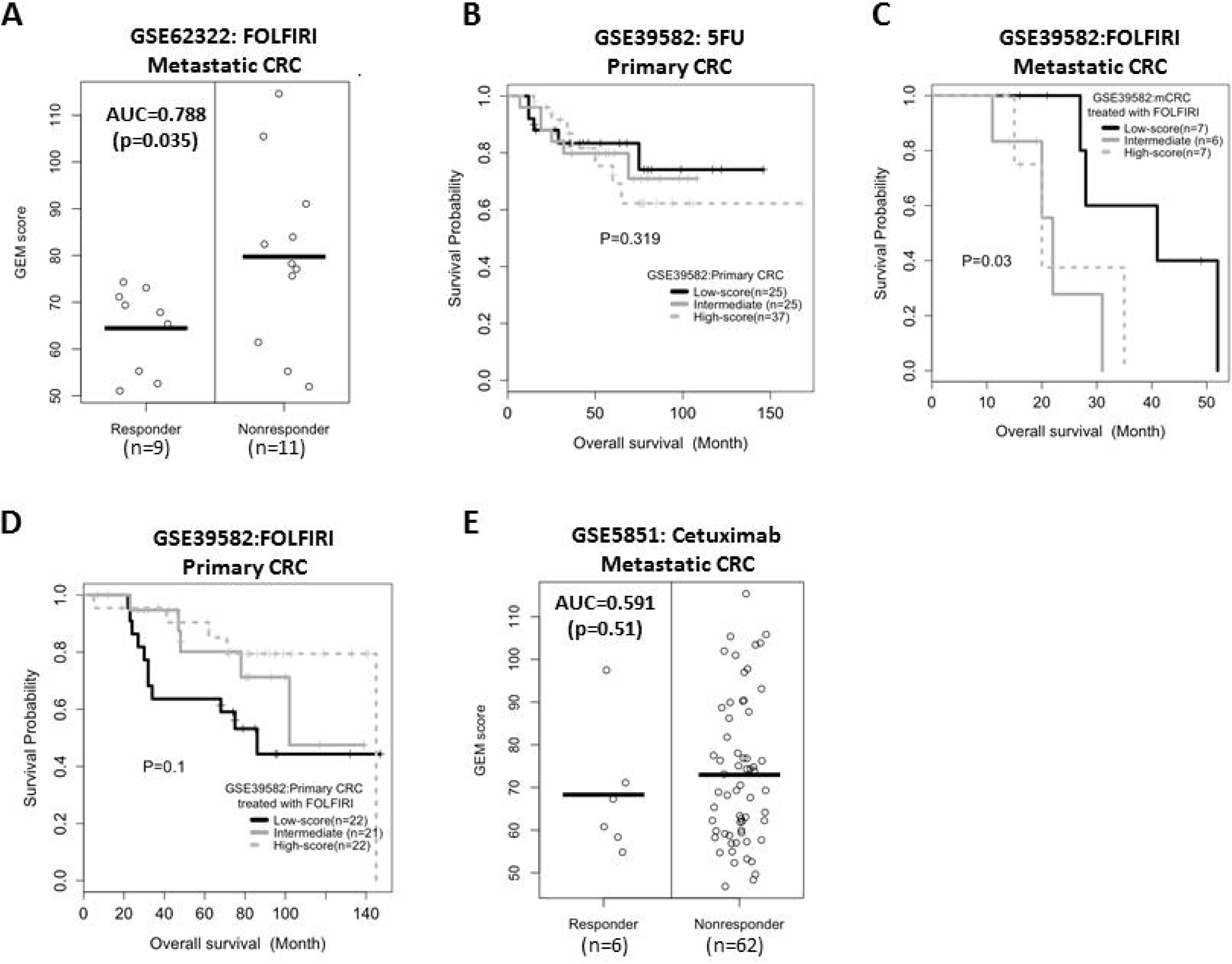
PDXGEM for 5FU response prediction in colorectal cancer patients. (a) Distribution of PDXGEM scores (Y-axis) between responsive and non-responsive patients after at a treatment with 5FU-based chemotherapy. (B-D) Kaplan-Meier curves of overall survival for the high (dotted gray), intermediate (gray), and low (black) score group in primary colorectal cancer patients receiving 5-FU monotherapy in GSE39581 (B), and metastatic CRC patients (C) and primary CRC patients (D) receiving FOLFIRI, Prediction scores were broken down at their tertiles. P-value was calculated using a survival trend test. (E) Distribution of PDXGEM scores (y-axis) of CRC patients who did not received 5-FU.

We next examined a prediction performance of our predictor in metastatic CRC patients from a prospective clinical trial for cetuximab monotherapy (GSE5851^40^) to assess whether the predictor predicts for 5FU-specific response. No significant difference in prediction scores was found (AUC=0.59, p=0.51), supporting that the PDXGEM predictor is predictive for 5FU-specific response (Figure 5E). +

### PDXGEM for Predicting Cetuximab Response in Colorectal Cancer Patients

Cetuximab is an antibody targeting the epidermal growth factor receptor (EGFR) and was approved for treating patients with metastatic CRC (mCRC) and patients with wild-type KRAS/NRAS/BRAF mCRC tumors are more likely to undergo disease control on cetuximab treatment. However, there is still an unmet need to identify effective biomarkers predicting treatment outcomes because a number of wild-type tumors especially in patients who do not have KRAS and/or BRAF mutations also remain unresponsive^41^. We thus applied PDXGEM to develop a predictive multi-gene signature of cetuximab response in patients with mCRC.

997 differentially expressed probesets were first selected by unpaired *t*-test analysis of nine sensitive and seven resistant PDXs after receiving cetuximab (p<0.05). We next screened out 670 biomarkers co-expressed concordantly across the PDXs and a cohort of mCRC patients (GSE14095^34^). An optimal RF predictor was then constructed based on 585 CCE biomarkers along with a strong correlation coefficient of 0.98 between prediction scores and observed percent of change in tumor volumes in the PDX training dataset.

We proceeded to an external validation using 68 mCRC patients who received cetuximab monotherapy in a phase 2 clinical trial (GSE5851^40^) and observed a significant difference in prediction scores between six responders and 62 non-responders (AUC=0.699, p=0.041; Figure 6A). When patients’ survival outcomes were analyzed in the same manner as described above, the high-score group showed worse progression-free survival with 6-months PFS rate of 3.7% than the low and intermediate score groups with 6-months PFS rate of 18.5% and 19.2%, respectively (Supplementary Fig. 5A; log-rank p=0.085).

**Figure 6.**
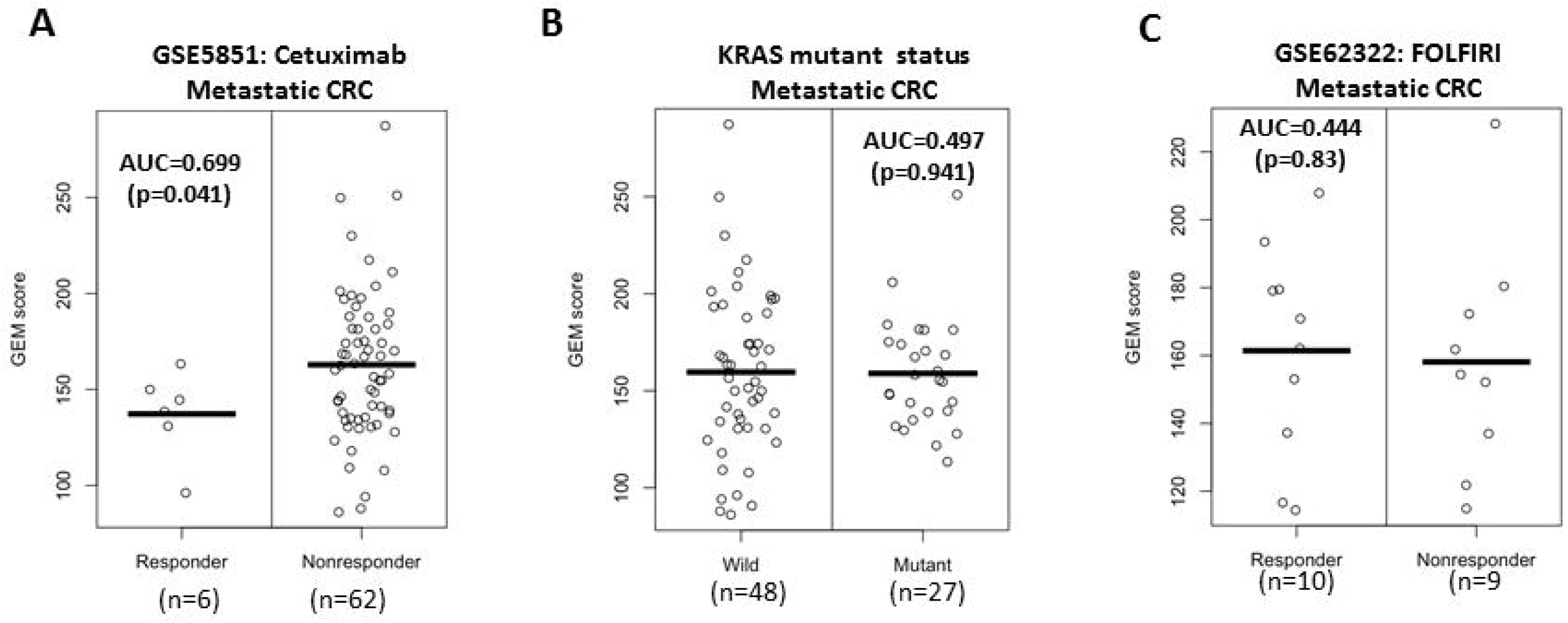
PDXGEM prediction for response to Cetuximab in metastatic colorectal cancer patient. (a) Distribution of PDXGEM scores (y-axis) between responsive and non-responsive CRC patients to cetuximab (b) PDXGEM scores stratified by Kras mutation status (c) Kaplan-Meier curves of overall survival in metastatic colorectal cancer patients who did not receive cetuximab.

Because cetuximab was already well known to be effective for metastatic CRC patients with wild type KRAS gene, we examined whether there is an association between the prediction scores and mutation status of KRAS gene in all GSE5851 patients. There was no significant difference in prediction \scores between patients with wild KRAS and those with mutant KRAS (p=0.941, Figure 6B). This result shows that the PDXGEM prediction score is independent of KRAS mutation status that is a well-known predictive biomarker in CRC patients. The PDXGEM score was able to screen further those with wild KRAS gene but non-responsive. (p=0.038; Supplementary Fig. 5B).

In determining whether the predictor has cetuximab specificity, we validated it on independent cohort of mCRC patients who did receive not cetuximab but FOLFIRI therapy (GSE62322^36^). No significant difference was seen in prediction scores between 9 responders and 10 non-responders (AUC=0.444; p=0.72; Figure 6C). This result implied that our predictor was predictive of response to cetuximab.

### PDXGEM Signature Predictive for Erlotinib Response in NSCLC Patients and Cell Clines

Erlotinib is an EGFR tyrosine kinase inhibitor and was approved for treatment of non-small cell lung cancer (NSCLC), but the overall therapeutic efficacy is minimal^42^. We attempted to construct a multi-gene signature for predicting response to the erlotinib by analyzing data on pretreatment gene expression profile and percent of changes in tumor volume s after erlotinib administration in eight NSCLC PDXs.

1,624 initial drug sensitivity biomarkers were screened using unpaired *t*-test that contrasts three PDXs with tumor shrinkage and five PDXs with tumor growth. 112 of them showed concordant co-expression patterns between the PDXs and a cohort of 150 NSCLC patients (GSE43580^43^). Finally, a 106-gene based RF predictor was built to predict a post-erlotinib treatment tumor volume change. Prediction scores of the RF predictor for the PDX training set was significantly correlated with the observed percent of changes in tumor volumes (r=0.974).

We first validated the predictor on gene expression data of *in vitro* erlotinib-treated NSCLC cell lines (GSE31625^44^; n=46). There was a significantly large difference in prediction scores between 18 erlotinib-sensitive cell lines and 28 erlotinib-resistant cell lines (AUC=0.708; p=0.006; see Figure 7A). We next validated the predictor on a prospective clinical trial cohort of 41 refractory NSCLC patients who received the first-line treatment with erlotinib in combination with bevacizumab(GSE37138^45^). We again observed a significant difference in prediction scores between 5 responders and 36 non-responders (AUC=0.689, p=0.016; Figure 7B). We next validated the predictor on 26 patients with relapsed or metastatic NSCLC who had EGFR mutation and received erlotinib at the second-line treatment setting (GSE33072^46^). Although our predictor yielded the highest prediction scores for two patients with the shortest progression-free survival times, there was no significant difference in PFS between high-score and low-score groups (see Figure 7C and Supplementary Figure 6A), implying that our prediction may not be predictive at the second line treatment setting.

**Figure 7.**
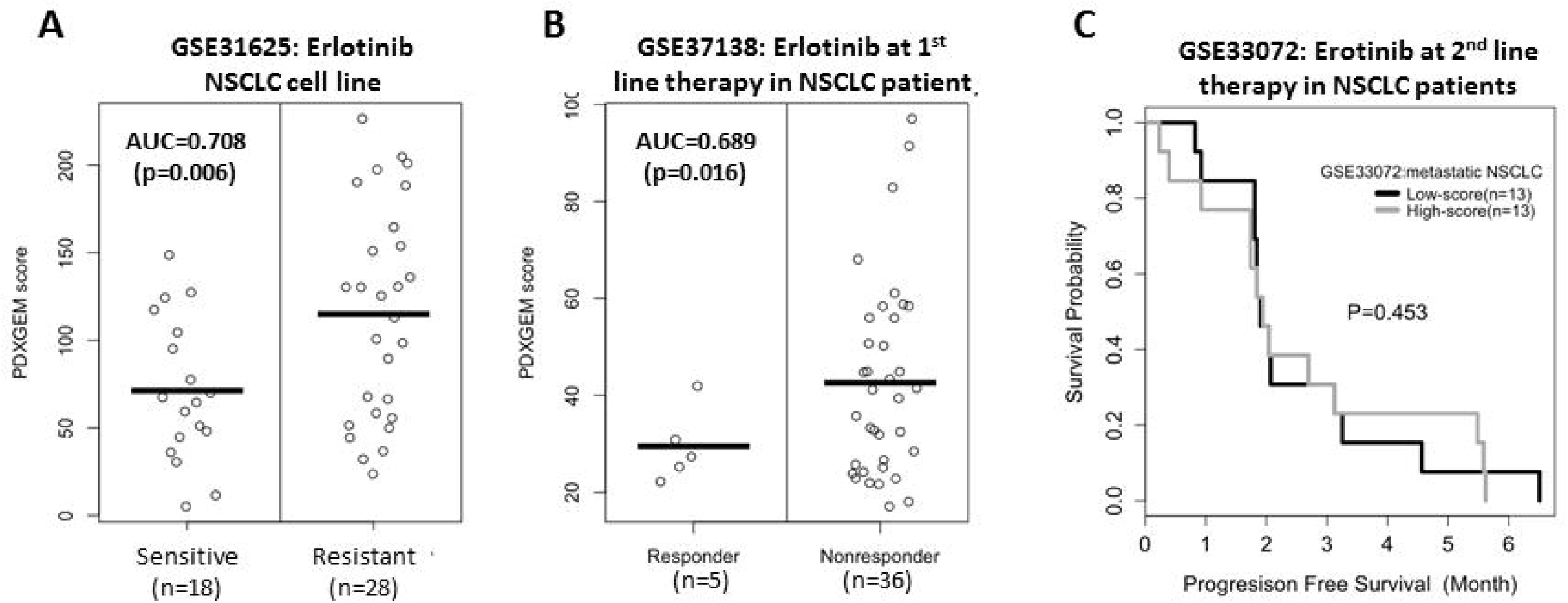
PDXGEM prediction for response to erlotinib in non-small cell lung cancer patient. (a) PDXGEM scores between erlotinib-sensitive and resistant NSCLC cell lines, and (b) between responsive and nonresponsive NSCLC patients after receiving erlotinib at the first line setting (c) progress-free survival curves in metastatic NSCLC patients who receive erlotinib as the 2^nd^ line treatment setting.

To examine erlotinib-specific response of the PDXGEM, we made a prediction in 20 NSCLC patients from The Cancer Genome Atlas (GSE68793) who did receive neither erlotinib nor other EGFR inhibitors such as gefitinib. There was no significant association of prediction scores with PFS and OS (Supplementary Figure 6B and 6C). Taken together, our validation results supported that our PDXGEM predictor could predict a response to erlotinib in chemo-naïve or EGFR wild-type NSCLC patients, but not at the second line treatment setting.

## Discussion

Predictive cancer biomarkers are necessary toward a personalized cancer treatment that will guide individual cancer patient to most likely effective therapy among a variety of anti-cancer drugs. Those studies that use PDX models for similar goals failed to provide exact equations of prediction models or especially software. Also, many approaches based on cancer cell line data has been developed including our recent studies, but again there is no publicly available software. We thus developed a statistical bioinformatics pipeline, PDXGEM, to facilitate building a multi-gene expression signature as a quantitative cancer biomarker for predicting a cancer patient’s response to a single anti-cancer drug on the basis of data on pretreatment gene expression profiles and post-treatment outcomes in preclinical PDX models. We demonstrated in this study that the PDXGEM can build predictive signatures for responses to chemotherapy as well as targeted therapy agents.

In this study, we devised CCEC statistics to quantify the degree of concordance of co-expression patterns between preclinical PDX tumors and cancer patient tumors because the PDX tumors can alter biological characteristics of their origin patient tumors to adapt new growth environments. Although drug sensitivity biomarkers obtained directly from a correlative analysis of data from preclinical PDX models could serve as predictive biomarkers by themselves, a subset of them with significant CCEC could induce a more translatable predictor yielding a better performance for predicting therapeutic outcomes in cancer patients as shown in our examples.

It is worthy to note that the PDXGEM does not use any patients’ outcome data during a prediction model development while various strategies for developing predictive gene signatures by analyzing data of preclinical models often uses patient’s outcome data for screening biomarkers or training a prediction model. The PDXGEM uses only pretreatment gene expression data of cancer patients at the concordant co-expression analysis step and it can therefore build gene signatures for anti-cancer drugs that are not approved for patients with a certain types of cancer. In addition, the PDXGEM uses a pretreatment gene expression data of cancer patients in CCEA analysis to improve GEMs based fully on PDX models. In the absence of available patient data, the PDXGEM build a prediction model by skipping the CCEA step.

Our PDXGEM applications demonstrated that highly predictive GEMs can be developed from a small cohort of PDX models. For example, the PDXGEM for erlotinib treatment response prediction utilizes only 8 PDX models, but its validation on NSCLC cancer cell lines as well as patients yielded statistically significant prediction performances. This is a highly desirable feature because there is frequently a limited resource of preclinical PDX models.

Indication of targeted therapy agents highly depends on status of their known target companion biomarkers in patient tumors. Our PDXGEM applications for Cetuximab can differentiate responders from non-responders within KRAS wild-type group based on PDXGEM scores. Usage of PDXGEM along with the known target biomarkers has a potential for improving clinical outcomes or quality of life in a population of such patients. In addition, the PDXGEM has a potential of being applicable to a recent breakthrough immunotherapy. Various immune-oncology studies recently began with creating and investigating PDX in mice with a human immune system^47^. Data collected from these humanized mice will open up a new application of PDXGEM for discovering and developing predictive biomarkers of response to the immunotherapy.

Many cancer drugs including drugs used in our examples are multi-indication drugs and can be used for treating more than one cancer type. For instance, paclitaxel is currently a standard chemotherapy drug for treating breast cancer as well as ovarian cancer. There is great interest in identifying a new treatment target of existing anti-cancer therapy agents. We recently introduced a drug repositioning approach, CONCORD, to translating predictive cancer biomarkers from one cancer type to another cancer type. The CONCORD analyzed data on gene expression and drug sensitivity data of a large panel of cancer cell lines with different types of cancer^13^. Similarly, there will be great interest in expanding PDXGEM to explore a potential of a drug for anti-cancer drug repositioning by testing prediction values of a predictive gene expression signature across multiple types of cancer because a larger resource of PDX panels that span multiple types of cancer is becoming publicly available.

There are clear limitations in developing the PDXGEM pipeline. The concordance of co-expression analysis result has dependency on a pretreatment gene expression data set of cancer patients that represents a cancer type of interest. Although we used as large gene expression dataset as possible, in terms of the number of patients and the coverage of histological cancer subtypes, it would be useful to have a more comprehensive gene expression data set for an individual cancer subtype by merging multiple independent gene expression datasets. In order to build a multi-gene expression model, we used the RF modeling algorithm to handle a larger number of gene biomarkers than a quite small sample size of the PDX data as a model training data. However, other statistical prediction modeling and machine learning algorithms such as penalized linear regression analysis and support vector machine could also be used build more accurately predictive models^48^. A majority of gene expression data we used was profiled on old-fashioned microarray platforms. Although we validated PDXGEM signature for gemcitabine on gene expression data profiled using RNAseq platform (ICGC cohort), it will be needed to validate its cross-platform prediction performance on other next generation sequencing dataset. Furthermore, a recent pharmacogenomics study of cancer cell lines reported that transcript-level expression profiled on RNAseq platform could lead more predictive biomarkers than a gene-level expression data. We also believe that application of PDXGEM to RNAseq transcriptional profiling data can lead to a predictive cancer biomarker with a better performance.

It will also be of utility to investigate whether PDXGEM can be extended to different molecular data such as genome-wide mutation data, proteomics, or metabolomics data. We believe that the mathematical framework of PDXGEM will be broadly applicable to these different molecular platforms. However, one may need to carefully examine if large reliable patient data resources are available and whether predictive therapeutic biomarkers can be obtained from such molecular data.

Lastly, we provided web-based PDXGEM application http://pdxgem.moffitt.org to share PDXGEM algorithm with the scientific community with hope that this tool will allow researchers to gain a better perspective of the drug targets and validation in a prospective study.

## Conclusions

Molecular profiles and drug activity data from PDX tumors can be used to develop highly predictive cancer biomarkers for predicting responses to anti-cancer drugs in cancer patients. The ultimate utility of these PDXGEM predictions should be assessed by a prospective study.

## Materials and Methods

### Gene Expression data and anti-cancer response data of PDX and Cancer Patient cohort

Data on gene expression profiles and post-treatment percent change in tumor volume of the Novartis PDX panel were obtained from Gene Expression Omnibus (GEO) repository (https://www.ncbi.nlm.nih.gov/geo/). Gene expression data or/and clinical outcome data of Cancer patients used for the concordant co-expression analysis (CCEA) or validation analysis werealsopubliclyavailableatGEO(http://www.ncbi.nlm.nih.gov/geo),ArrayExpress (https://www.ebi.ac.uk/arrayexpress/),and International Cancer Genome Consortium(https://dcc.icgc.org/repository). A descriptive summary and accession ID of all the data can be found in **Supplementary Table 1.**

### *In Vivo* PDX-based Drug Sensitivity Biomarker Discovery

We discovered genes whose expression levels were significantly associated with each drug’s *in vivo* activities on PDX tumors for the target cancer type of a given anti-cancer drug. The drug activity was calculated as a percent change in PDX tumor volumes (100 x (post-treatment tumor volume – pretreatment tumor volume) / pretreatment tumor volume). A negative drug activity value for a PDX thus means a tumor shrinkage while a positive drug activity value represents a tumor growth. The drug activity data and pretreatment gene expression profiling data of the PDX models were analyzed to screen the most accurate drug sensitivity biomarkers for the drug. The basic unit of biomarker is an individual probeset in the microarray data of the PDXs. Drug sensitivity biomarkers were selected using the unpaired two sample *t*-test that quantifies differential gene expression levels between PDXs with shrunken tumors and those with grown tumors. When a sample size in one of the two PDX groups was less than three and a variation of tumor volume changes was small near zero, a correlation analysis of gene expression levels with the percent change of tumor volumes was used for screening the initial drug sensitivity biomarkers. For both t-test and correlation analyses, all statistical tests were two-sided and the false discovery rate was controlled to be less than 0.05 to correct for multiple comparisons. When no significant genes were found due mainly to a small sample size of available PDXs, we controlled a less conservative nominal type I error rate of 0.05 to identify initial drug sensitivity biomarkers.

### Concordant Co-Expression Analysis (CCEA)

Since PDX tumors can alter biological characteristics of their origin patient tumors to adapt new growth environments, not all the drug sensitivity genes screened from an analysis of data from the PDX tumors may be predictive of response of cancer patients. To explicitly take into account such a biological difference, we selected genes with concordant co-expression patterns between the PDX tumors and cancer patient tumors. To quantify the degree of concordance of each gene’s co-expression relationships, we calculated the concordance co-expression coefficient (CCEC) for each gene *g* as follows. Using expression data within each of the two cancer systems separately, we first constructed two correlation matrices (of dimension *n*×*n*) for *n* drug-sensitivity biomarkers. The two correlation matrices, e.g. one for the PDX tumor set and another for the pretreatment cancer patient tumor set, both were evaluated as *U = [U*_*ij*_]_*n*×*n*_ and *V = [V*_*ij*_]_*n*×*n*_, where *U*_*ij*_ and *V*_*ij*_ are the correlation coefficients between probeset *i* and *j* in the PDXs and the patient set, respectively. Then, the CCEC for the gene *g, c(g)*, is derived as

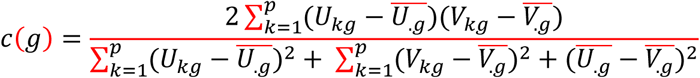

where 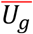 and 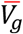 are the mean correlation coefficients of the *j-*th column correlation coefficient vectors for the PDX and patient cohort. The CCEC *c(g)* quantifies the degree of agreement between two correlation coefficient vectors *Ug* and *Vg* using the concordance correlation coefficient^49^. Therefore, *c(g)* reflects in the example of paclitaxel the degree of concordance between the breast cancer PDX panel and BR-251 cancer patient cohort for expression relationships of probe *g* with other *n-1* probesets. If *c(g)* has a statistically significant positive value under false discovery rate of 0.05, probeset *g* is selected as a significant CCE biomarker. Note that since the probeset *g* was initially selected among *n* drug-sensitivity biomarkers, it retained a significant association with drug sensitivity.

### PDXGEM Modeling and Evaluation

A multi-gene expression model for predicting each drug’s response was built using gene expression data and drug activity data of the PDX panel that was used in the above drug sensitivity biomarker discovery. The drug activity data and gene expression data of the PDX model for all CCE biomarkers, defined as drug sensitivity genes with statistically significant CCEC, formed the model training data. A random forest classification and regression analysis was performed after completing a gene-wise standardization of the model training data. The prediction performance of the resultant RF predictor was first evaluated by calculating a correlation coefficient between the observed and predicted tumor volume changes in the PDX mouse models. When there was a significant correlation relationship, the RF predictor was validated on gene expression data and post-treatment clinical outcome data of cancer patient cohorts that were completely independent of the biomarker discovery and the prediction model development.

### PDXGEM Prediction and Validation

To validate the prediction performance of each drug’s final prediction model, we applied it to historical cancer patient cohorts that were completely independent of drug sensitivity biomarker discovery and prediction model development procedures. The performance of each drug’s PDXGEM prediction was then assessed in a prospective manner. For cancer patient cohorts with binary response outcome data, we compared prediction scores between responsive and non-responsive patient groups by unpaired two-sample *t*-test. The area under the receiver operating characteristic curve (AUC) was also calculated to summarize an overall prediction performance of the prediction model. For cancer patient cohorts with survival outcome data, survival distributions were compared between their prediction score strata by Kaplan-Meier survival analysis and a log-rank test. Multivariable Cox proportional hazard regression analysis was also used to examine an association of raw continuous prediction scores with survival outcomes.

### Gene Ontology Analysis

To understand any potent gene functional behaviors and mechanisms with which the multi-gene expression model could predict patients’ responses to the anticancer drug of interest, we selected CCE biomarkers showing a positive value of variable importance in the RF analysis. In brief, the variable importance is a model selection measure of the RF analysis by summarizing a difference in prediction accuracy between two RF predictors with and without individual biomarker as a model selection measure. Finally, a web-based Enrichr tool was used for the GO analysis of the selected CCE biomarkers^50^.

## List Abbreviation

AUC: Area under Receiver Operating Characteristic Curve
CCEA: Concordance Co-Expression Analysis
CCEC: Concordance Co-Expression Coefficient
CCR: Colorectal Cancer
FAC: fluorouracil, doxorubicin, and cyclophosphamide
5FU: 5-fluoracil
FOLFIRI: Fluoriacil, leucovorin, and Irinotecan
GEM: Gene Expression Model
GO: Gene Ontology
ICGC: International Cancer Genome Consortium
NSCLC: Non-Small Cell Lung Cancer
PDX: Patient-Derived Xenograft
ROC: Receiver operating Characteristic curve

## Availability of data and materials

The datasets analysed during the current study are available with data accession IDs presented in the manuscript in the Gene Expression Omnibus repository, https://www.ncbi.nlm.nih.gov, and ArrayExpress repository, https://www.ebi.ac.uk/arrayexpress. Data access ids are presented in the manuscript and supplementary Table 1.

## Authors’ Contributions

Conception and design: YK and DK; collection and assembly of data: YK, BC; data analysis and interpretation: YK, MK and DK; software development: RC; manuscript writing: YK, BC, RC, MK, and DK; final approval of manuscript: All authors read and approved for publication.

## Acknowledgements

This study was supported by the Shared Resources at the H. Lee Moffitt Cancer Center and Research Institute, an NCI designated Comprehensive Cancer Center (P30-CA076292).

