## Supplementary figures and images for "PDXGEM: Patient-Derived Tumor Xenograft based Gene Expression Model for Predicting Clinical Response to Anticancer Therapy in Cancer Patients"

### Supplementary Figure 1

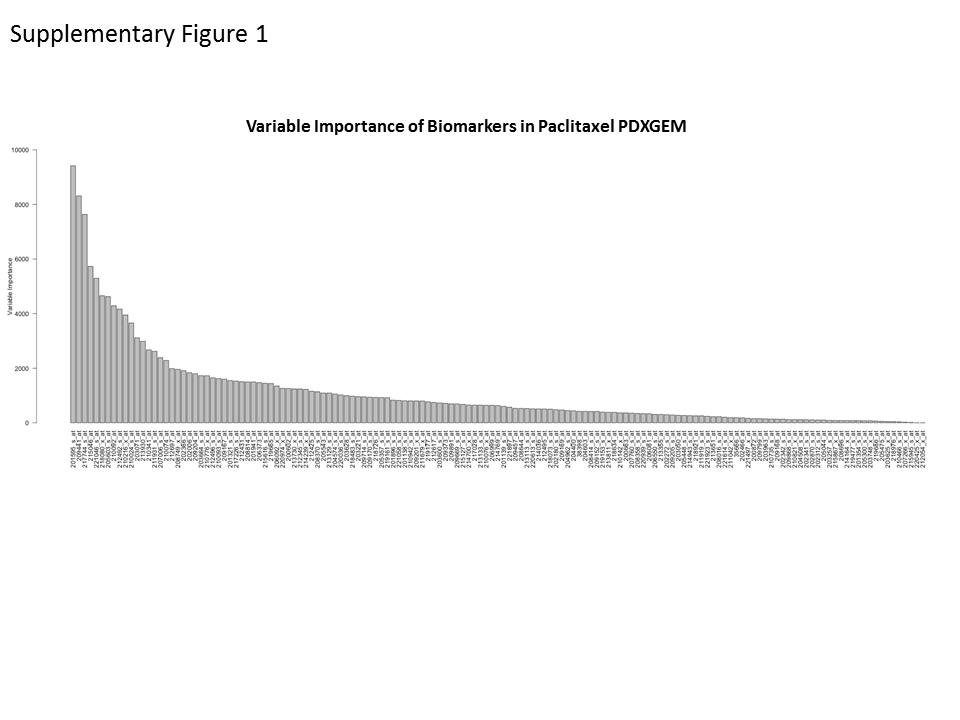

### Supplementary Figure 2

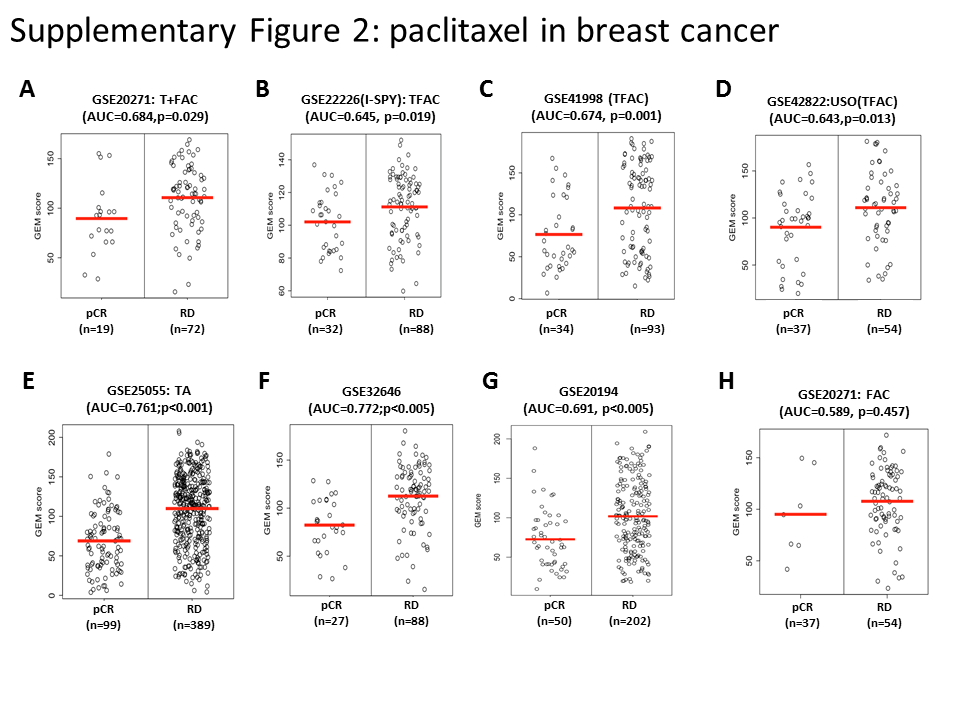

### Supplementary Figure 3

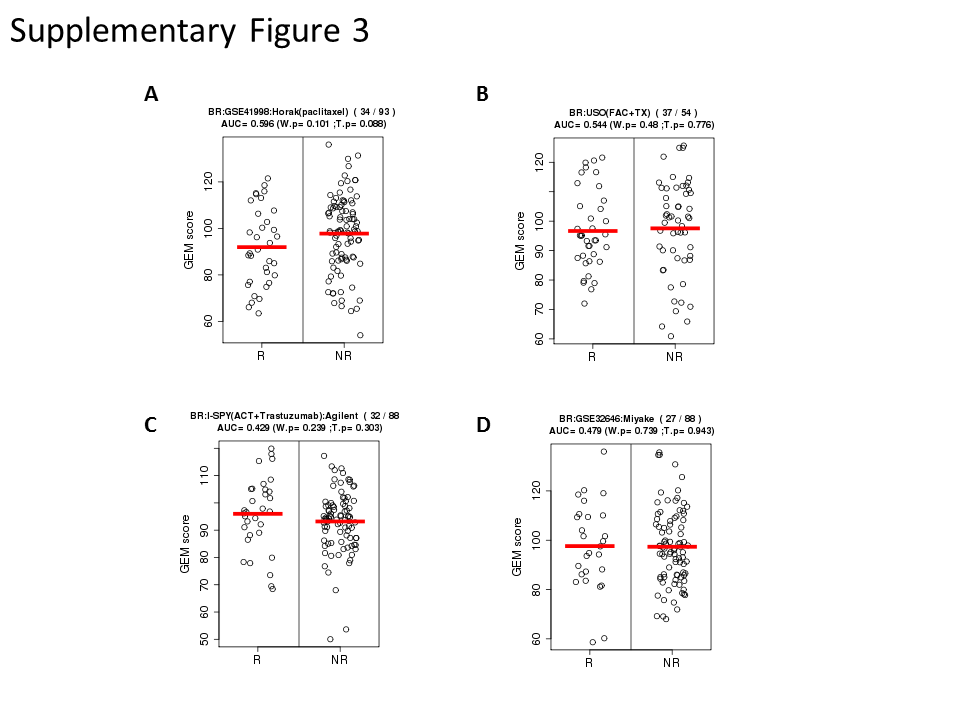

### Supplementary Figure 4

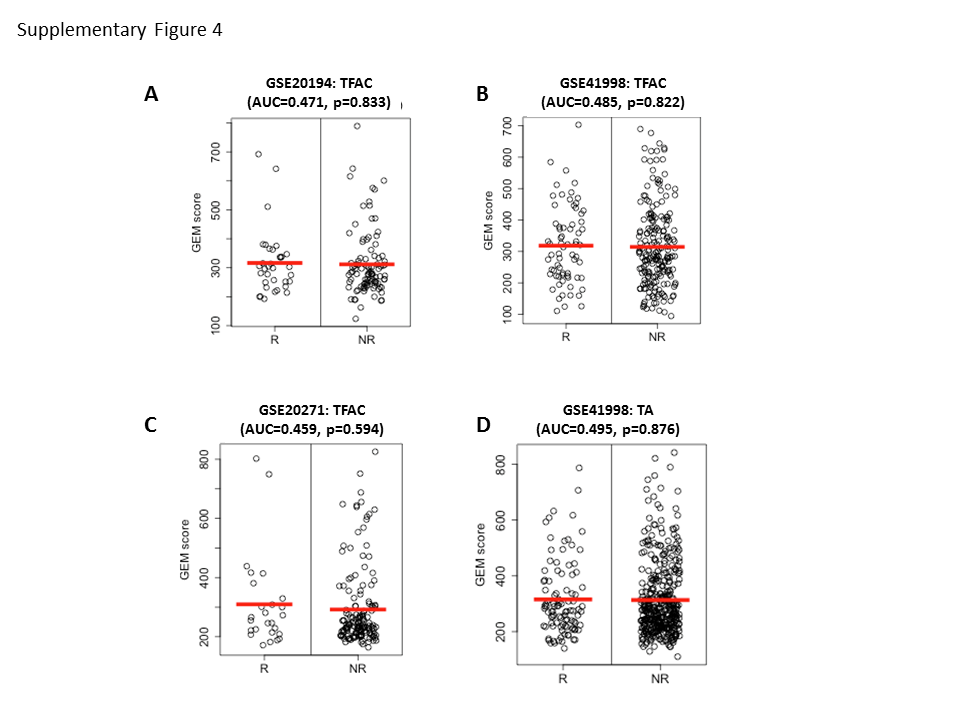

### Supplementary Figure 5

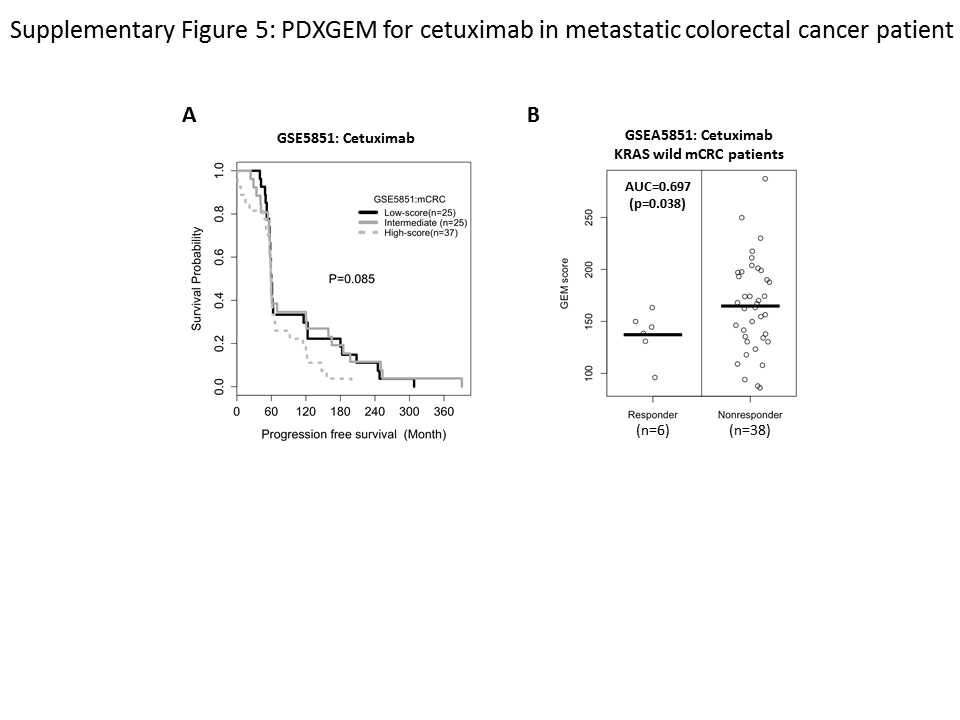

### Supplementary Figure 6

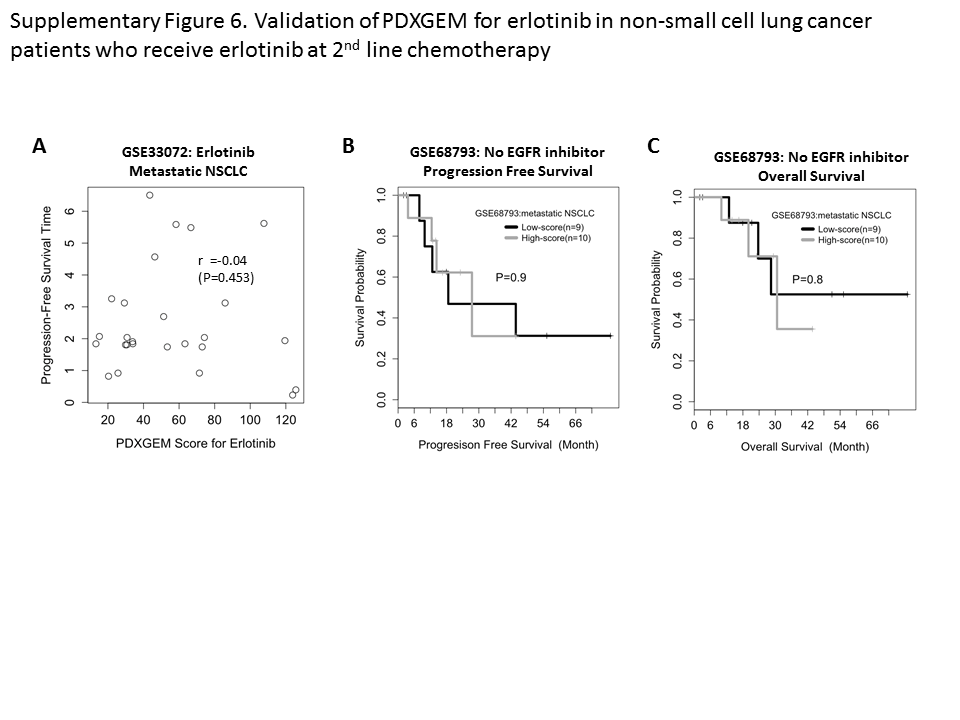
