## Supplementary Table 1 for "PDXGEM: Patient-Derived Tumor Xenograft based Gene Expression Model for Predicting Clinical Response to Anticancer Therapy in Cancer Patients"

**Supplementary Table 1.** The list of gene expression and anti-cancer drug response data sets

| Cancer type | GEO access ID or website | Study type/ name | Gene Expression profiling platform | Drug response | Number of patients | Anti-cancer Drugs | Role of data |
|---|---|---|---|---|---|---|---|
| PDX | GSE78806 | Novartis PDX Panel | U133A | % change in tumor volume | | | Biomarker Discovery& Model Training set |
| Breast Cancer | GSE3494 (Miller et al.) | retrospective tumor bank study | U133A | | 251 | - | CCEA set |
| | GSE20271 (Tabchy et al.) | Clinical trial | U133A | pCR | 91 TFAC | T+FAC or FAC | validation |
| | GSE22226 (Esserman et al.) | Clinical trial (I-SPY) | Agilent 4x44k | pCR | 120 TFAC | | validation |
| | GSE41998 (Horak et al.) | Clinical trial (NCT00455533) | U133A 2.0 | pCR | 127 | T+AC | validation |
| | GSE25065 (Hatzis et al) | validation set in a prospective biomarker study | U133A | pCR | 498 | T+A | |
| | GSE42822 (Shen et al) | Clinical trial (USO 02-103) | U133A | pCR | 91 TFAC | Trastuzumab + TFAC or TFAC alone | Validation |
| | GSE32646 (Miyake et al) | ER-negative breast cancer study in Japan | U133+2 | pCR | 115 | TFEC | validation |
| | GSE20194 (Shi et al.) | MAQC-II study | U133A | pCR | 278 | TFAC | validation |
| pancreatic ductal adenocarcinoma | GSE15471 (Badea et al) | Non-clinical trial | U133+2 | | 78 | | CCEA |
| | GSE57495 (Chen et al.) | Non-clinical trial | Affymetrix HuRSTA | OS | 63 | Gemcitabine | validation |
| | E-MEXP-2780* (Winter et al.) | Non-clinical trial | U133+2 | OS | 30 | N/A | Validation |
| | ICGC | ICGC | RNASeq | OS | 96 | N/A | Validation |

| Cancer type | GEO access ID or website | Study type/ name | Gene Expression profiling platform | Drug response | Number of patients | Anti-cancer Drugs | Role of data |
|---|---|---|---|---|---|---|---|
| | GSE17891 (Collison et al) | tumor bank study at UCSF | U133+2 | OS | | N/A | validation |
| Colorectal Cancer | GSE14095 (Watanabe et al.) | Non-clinical trial | U133+2 | | 189 | | CCEA |
| | GSE62322 (Del et al.) | Non-clinical trial | U133A &U133B | pCR | 114 | FOLFIRI | validation |
| | GSE39582 (Marisa et al.) | Non-clinical trial | U133+2 | OS | 585 | FOLFIRI or FOLFOX | validation |
| | GSE5851 (Khambata-Ford et al.); | prospective randomized trial | U133A2.0 | pCR | 80 | Cetuximab monotherapy | Validation |
| Non-small cell lung cancer | GSE43580 (Tarca et al.) | Non-clinical trial | U133+2 | | 150 | | CCEA |
| | GSE31625[$] (Balko et al.) | Cancer cell line Drug screening | U133A | Erlotinib-sensitivity | 18 / 28 | Erlotinib | validation |
| | GSE37138 (Baty et al.) | Prospective clinical trial (SAKK 19/05) | HuEx-1.0 ST | pCR | 117 | Erlotinib+ Bevacizumab | validation |
| | GSE33072 (Byers et al.) | Clinical trial (BATTLE) | HuEx-1.0 ST | PFS | | Erlotinib at 2$^{nd}$ line therapy | validation |
| | GSE68793 (TCGA) | TCGA Consortium study | U133A | PFS OS | 135 | Non EGFR-inhibitors | validation |

**TFAC: paclitaxel, 5-FU, Adriamycin and Cyclophosphamide;**
**TFEC: paclitaxel, 5-FU, Epirubicin and Cyclophosphamide**
**FOLFIRI: folinic acid, 5FU, and irinotecan;**
**FOLFOX: folinic acid, 5F and oxaliplatin**
**pCR** : pathologic complete response, **OS** : overall survival, **PFS** : progression free survival
[*] : Access ID of gene expression data at ArrayExpress
[$:] Cancer cell line study
N/A: not available
